# Genetic Conservation of SARS-CoV-2 RNA Replication Complex in Globally Circulating Isolates and Recently Emerged Variants from Humans and Minks Suggests Minimal Pre-Existing Resistance to Remdesivir

**DOI:** 10.1101/2020.12.19.423600

**Authors:** Ross Martin, Jiani Li, Aiyippa Parvangada, Jason Perry, Tomas Cihlar, Hongmei Mo, Danielle Porter, Evguenia Svarovskaia

**Author notes:** Corresponding author: Ross Martin Gilead Sciences, Inc., 333 Lakeside Dr, Foster City, CA.

## Abstract

Remdesivir (RDV) exhibits potent antiviral activity against SARS-CoV-2 and is currently the only drug approved for the treatment of COVID-19. However, little is currently known about the potential for pre-existing resistance to RDV and the possibility of SARS-CoV-2 genetic diversification that might impact RDV efficacy as the virus continue to spread globally. In this study, >90,000 SARS-CoV-2 sequences from globally circulating clinical isolates, including sequences from recently emerged United Kingdom and South Africa variants, and >300 from mink isolates were analyzed for genetic diversity in the RNA replication complex (nsp7, nsp8, nsp10, nsp12, nsp13, and nsp14) with a focus on the RNA-dependent RNA polymerase (nsp12), the molecular target of RDV. Overall, low genetic variation was observed with only 12 amino acid substitutions present in the entire RNA replication complex in ≥0.5% of analyzed sequences with the highest overall frequency (82.2%) observed for nsp12 P323L that consistently increased over time. Low sequence variation in the RNA replication complex was also observed among the mink isolates. Importantly, the coronavirus Nsp12 mutations previously selected in vitro in the presence of RDV were identified in only 2 isolates (0.002%) within all the analyzed sequences. In addition, among the sequence variants observed in ≥0.5% clinical isolates, including P323L, none were located near the established polymerase active site or sites critical for the RDV mechanism of inhibition. In summary, the low diversity and high genetic stability of the RNA replication complex observed over time and in the recently emerged SARS-CoV-2 variants suggests a minimal global risk of pre-existing SARS-CoV-2 resistance to RDV.

## 1. Introduction

Over the past two decades, emerging pathogenic coronaviruses (CoVs) that can cause life-threatening diseases in humans and animals have been identified, namely severe acute respiratory syndrome coronavirus (SARS-CoV), Middle Eastern respiratory syndrome coronavirus (MERS-CoV), and most recently SARS-CoV-2, the causative agent of coronavirus disease 2019 (COVID-19) [1]. A critical step in the coronavirus lifecycle, including SARS-CoV-2, involves the replication of the viral RNA. This replication is a complex process that involves full genome replication, subgenomic RNA transcription, proofreading, and mRNA capping [2] The viral RNA replication complex of SARS-CoV-2 is composed of multiple proteins including an RNA dependent RNA polymerase (nsp12), a zinc-binding helicase (nsp13), and a proofreading exonuclease (nsp14) [2]. RNA polymerization depends on the main polymerase catalytic subunit nsp12, which is bound to the accessory subunits nsp7 and nsp8 [3]. Nsp13 is a helicase which has been shown to associate with the nsp12/nsp7/nsp8 complex [4] and likely functions as a translocase in the complex, enabling RNA backtracking, which may be critical to both proofreading and transcription. Nsp14 is responsible for proofreading at its exonuclease active site, as well as a key step in mRNA capping at its *N7*-methyltransferase site [3]. Exonuclease activity is enhanced by nsp14’s association with nsp10, but how these two proteins interact with the complex is still not known. The exonuclease activity in SARS-CoV-2 and other coronaviruses results in a lower mutation rate and the ability to maintain genetic stability of their large RNA genome as compared to other RNA viruses

Remdesivir (RDV; GS-5734) is a single diastereomer monophosphoramidate prodrug of a nucleoside analog that is intracellularly metabolized into an analog of adenosine triphosphate which inhibits viral RNA polymerases. RDV exhibits a potent in vitro broad spectrum activity against members of the coronaviruses, filoviruses, and paramyxoviruses with EC50 values ranging from 0.0035 to 0.75 μM [5–7]. Importantly, RDV has also shown potent antiviral activity against SARS-CoV-2 in vitro with EC50 values as low as 0.01 uM in primary human airway epithelial cells [8] and 0.13 uM in human lung cell lines [9]. RDV has also demonstrated in vivo therapeutic efficacy against SARS-CoV-2 in rhesus macaques [10] and in vivo prophylactic and therapeutic efficacy against SARS-CoV and MERS-CoV infection in mice and MERS-CoV infection in rhesus monkeys [5, 11, 12]. Importantly, results from the Phase 3 randomized, double-blind, placebo-controlled Adaptive Covid-19 Treatment Trial (ACTT-1) showed that a 10-day treatment with RDV significantly shortened the time to recovery compared to control placebo group in adults who were hospitalized with COVID-19 [13]. In addition, two randomized open-label studies have demonstrated efficacy of shorter 5-day RDV treatment in hospitalized COVID-19 patients [14, 15]. RDV is currently the only drug approved by the US Food and Drug Administration for the treatment of COVID-19.

Biochemically, remdesivir triphosphate (RDV-TP) has been shown to inhibit SARS-CoV-2 RNA replication by two mechanisms that target nsp12 [16, 17]. The first opportunity to inhibit RNA synthesis is via delayed chain termination. RDV-TP has been observed to be incorporated by the polymerase into the nascent RNA strand as a mimic of ATP. Synthesis continues with the addition of three more nucleotides, before the process is halted. This pause in replication has been linked to a steric clash between the 1’-cyano of RDV and residue S861 of nsp12 [17]. This was confirmed by an engineered S861G mutation, which eliminated the delayed chain termination of RDV-TP. It has also been confirmed by recent structural studies [18]. A second opportunity to inhibit RNA synthesis was revealed for situations in which delayed chain termination does not occur and RDV-TP is stably incorporated into the RNA. In this case, with the new negative sense RNA copy now serving as a template, the incorporated RDV causes a misalignment when it reaches the nsp12 active site, due to a steric clash between the 1’-cyano and the protein at residue A558 [16]. As a result, the complementary UTP is inefficiently incorporated and further synthesis is compromised, leading to template-dependent inhibition.

The in vitro development of resistance to RDV in CoVs has been assessed by cell culture passaging of murine hepatitis virus (MHV) in the presence of GS-441524, the parent nucleoside of RDV. After 23 passages, two mutations were selected in the nsp12 polymerase at residues F476L and V553L [19]. Compared with wild-type virus, recombinant MHV containing the F476L and V553L single mutation showed 2.4-fold and 5-fold reduced susceptibility to RDV, respectively, while the double mutant conferred 5.6-fold reduced susceptibility to RDV in vitro. The mutant viruses were unable to compete with wild-type virus in coinfection experiments in the absence of RDV, demonstrating a viral fitness cost associated with these mutations. Corresponding mutations also confer similar reduction in RDV susceptibility when genetically transferred to recombinant SARS-CoV [19]. The F476 and V553 residues in MHV correspond to F480 and V557, respectively, in SARS-CoV-2. The substitution V557L was introduced in biochemical studies, where it was shown that the mutation impacted the template-dependent inhibition of RDV, but not RDV-TP incorporation or delayed chain termination [16]. Based on high conservation of the RdRp across coronaviruses it is likely that the resistance pathways will be similar; however, this needs to be experimentally determined.

In early studies, minimal genetic differences were observed among SARS-CoV-2 isolates from infected patients, with more than 99.9% sequence identity [20]. Further investigation is needed to monitor the emergence of polymorphisms as SARS-CoV-2 rapidly spreads in the human population worldwide and to other species. In December 2020, a new emergent SARS-CoV-2 variant was described as rapidly spreading across the United Kingdom [21]. The B.1.1.7 variant contained a signature of spike amino acid changes including H69/V70 deletion, Y144 deletion, N501Y, A570D, D614G, P681H, T716I, S982A, and D1118H. Another SARS-CoV-2 variant, B.1.351 commonly referred to as 501Y.V2, was described to be spreading quickly across South Africa [22]. The B.1.351 variant also contains a signature of spike amino acid changes including K417N, E484K, and N501Y. In addition, the transmission of SARS-CoV-2 between humans and minks has been reported near mink farms in Denmark and the Netherlands [23]. These events raise concerns of possible further diversification of the SARS-CoV-2 genomes and the potential impact on the efficacy of therapeutics and vaccines.

In this report, we analyzed amino acid substitutions in all critical parts of the RNA replication complex of SARS-CoV-2 in geographically diverse human clinical isolates including the recent emergent United Kingdom and South Africa variants and isolates from minks using publicly available sequences to assess global genetic diversification and its potential impact on the susceptibility of circulating viruses to RDV.

## 2. METHODS

### 2.1 SARS-CoV-2 Sequence Analyses

SARS-CoV-2 full genome sequences from clinical isolates and minks collected from December 2019 through early September 2020 were obtained from GISAID EpiCov database [24]. Sequences were aligned to the Wuhan-Hu-1 viral isolate sequence (NC_045512) using the Mafft Sequence Aligner v7.394 [25]. Nucleotide and amino acid changes across the SARS-CoV-2 genome were tabulated. Sequences (N=3,590 and N=5 clinical and mink isolate sequences, respectively) were excluded that contained ambiguous bases within RdRp (nsp12). A total of N=92,334 clinical isolate sequences and N=333 mink sequences were included in the analysis. The nsp12 nucleotide sequences (13442-16236) were adjusted for the ribosomal slippage at nucleotide position 13468 and translated for amino acid analysis. For nucleotide and amino acid analysis, if indels spanned multiple consecutive positions, only one change was counted. Ambiguous nucleotides (N) were ignored in counts of nucleotide changes. A linear least-squares regression was used to find trends in frequencies of amino acid substitutions over time (per month).

Additional SARS-CoV-2 sequences containing Spike N501Y mutation were downloaded on December 21, 2020. These sequences were separated into B.1.1.7 variant if they also contained spike H69/V70 deletion, Y144 deletion, A570D, D614G, P681H, T716I, S982A, and D1118H, and separated into B.1.351 variant if they also contained spike N417N and E484K. Based on this criteria, N=1987 B.1.1.7 and N=311 B.1.351 sequences were identified for analysis.

### 2.2 Pan-Coronavirus Nsp12 Sequence Analysis

To evaluate amino acid conservation of RdRp (Nsp12) in coronavirus subtypes, full genome sequences were aligned using Mafft Sequence Aligner v7.394 [25]. The nucleotide sequences were adjusted for the ribosomal slippage at nucleotide position 13468, relative to SARS-CoV-2, and translated for amino acid analysis. Coronavirus sequences included in the analysis were NC_038861 (TGEV), DQ010921 (FIPV), JX503060 (229E), AY567487 (NL63), MK841495 (PEDV), AY903460 (OC43), NC_003045 (BCV), DQ415904 (HKU1), NC_001846 (MHV-A59), MG772933 (Bat-SL-ZC45), MG772934 (Bat-SL-ZXC21), MN908947 (SARS-CoV-2), AY278741 (SARS-CoV-1), JX869059 (MERS), NC_009021 (Bat-HUK9-1), NC_001451 (IBV), NC_010646 (SW1), NC_011547 (HKU11), NC_016992 (HKU17). For investigation of sequence diversity within other coronaviruses, N=647 isolate sequences from human coronavirus infections were downloaded from NCBI Virus database (https://www.ncbi.nlm.nih.gov/labs/virus).

## 3. Results

### 3.1 Sequence Analyses of SARS-CoV-2 from Human Clinical and Mink Isolates Collected December 2019 to September 2020

To assess the genetic variation of nsp12 and other proteins involved in RNA replication, publicly available SARS-CoV-2 sequences from the ongoing pandemic were downloaded. A total of N=92,334 full genome sequences from human clinical isolates and N=333 isolates from minks were selected and aligned to the reference strain Wuhan-Hu-1 viral isolate (NC_ 045512). To note, the USA-WA-1 viral isolate (MN985325), to which most RDV antiviral activity testing was previously performed, contains no amino acid substitutions in the RNA replication complex as compared to the Wuhan-Hu-1 viral isolate. The number of sequences obtained, together with time and geographical location of sample collections are summarized in Table 1. The clinical isolate sequences were obtained from 109 countries from December 2019 up to September 2020. The mink sequences were obtained from Netherlands and Denmark from April to September 2020.

**Table 1.** Countries and Collection Dates of Human Clinical and Mink Isolates Analyzed.

| Host | Country | December 2019 | January 2020 | February 2020 | March 2020 | April 2020 | May 2020 | June 2020 | July 2020 | August 2020 | September 2020 | No Date | Total |
| --- | --- | --- | --- | --- | --- | --- | --- | --- | --- | --- | --- | --- | --- |
| Human | **Total** | **18** | **453** | **1,134** | **29,650** | **27,118** | **11,834** | **7,927** | **7,932** | **5,026** | **301** | **941** | **92,334** |
|  | United Kingdom |  | 15 | 108 | 9,318 | 14,653 | 4,823 | 1,357 | 1,060 | 2,817 | 122 | 506 | 34,779 |
|  | USA |  | 14 | 106 | 6,246 | 5,438 | 3,705 | 3,773 | 1,286 | 492 | 156 | 22 | 21,238 |
|  | Australia |  | 16 | 19 | 1,618 | 576 | 193 | 502 | 3,389 | 141 | 7 | 144 | 6,605 |
|  | Spain |  |  | 21 | 1,834 | 851 | 138 | 62 | 44 | 41 |  | 2 | 2,993 |
|  | India |  | 2 |  | 129 | 360 | 784 | 676 | 95 | 138 | 2 | 28 | 2,214 |
|  | Netherlands |  | 3 | 14 | 770 | 552 | 342 | 145 | 74 | 112 |  | 195 | 2,207 |
|  | Canada |  | 4 | 7 | 1,143 | 526 | 55 | 5 | 1 |  |  | 7 | 1,748 |
|  | Switzerland |  |  | 26 | 736 | 157 | 49 | 66 | 273 | 293 |  |  | 1,600 |
|  | Portugal |  |  |  | 1,079 | 199 | 63 | 57 |  |  |  |  | 1,398 |
|  | South Africa |  |  |  | 32 | 79 | 45 | 210 | 735 | 154 |  |  | 1,255 |
|  | Belgium |  |  | 2 | 555 | 270 | 73 | 14 | 96 | 67 |  |  | 1,077 |
|  | Ireland |  |  |  | 20 | 155 | 193 | 7 | 134 | 510 |  |  | 1,019 |
|  | Singapore |  | 11 | 39 | 257 | 320 | 78 | 91 | 94 | 45 | 14 | 1 | 950 |
|  | China | 18 | 308 | 334 | 186 | 4 | 2 | 3 | 51 | 3 |  |  | 909 |
|  | South Korea |  | 8 | 102 | 96 | 13 | 146 | 229 | 176 | 2 |  |  | 772 |
|  | Denmark |  |  | 2 | 568 | 79 | 2 |  |  |  |  |  | 651 |
|  | Japan |  | 10 | 123 | 320 | 151 | 21 | 1 |  |  |  |  | 626 |
|  | Sweden |  | 1 | 6 | 346 | 117 | 75 | 35 | 22 | 12 |  |  | 614 |
|  | Italy |  | 4 | 74 | 389 | 76 | 37 | 6 | 13 | 1 |  | 10 | 610 |
|  | Saudi Arabia |  |  | 16 | 85 | 295 | 76 | 90 |  | 8 |  |  | 570 |
|  | France |  | 8 | 16 | 430 | 95 |  | 11 |  |  |  |  | 560 |
|  | Iceland |  |  | 1 | 552 |  |  |  |  |  |  |  | 553 |
|  | Brazil |  |  | 5 | 136 | 318 | 40 | 4 | 2 | 1 |  |  | 506 |
| Mink | **Total** | **0** | **0** | **0** | **0** | **25** | **22** | **53** | **46** | **54** | **114** | **19** | **333** |
|  | Netherlands |  |  |  |  | 25 | 22 | 41 | 46 | 43 | 50 | 19 | 246 |
|  | Denmark |  |  |  |  |  |  | 12 |  | 11 | 64 |  | 87 |

### 3.2 Genetic Changes in SARS-CoV-2 Genome Since the Start of the Pandemic

The number of nucleotide changes relative to the reference strain Wuhan-Hu-1 (NC_ 045512) across the full genome and counts of amino acid substitutions across all ORFs for each human clinical isolate was calculated. Among the 92,334 analyzed SARS-CoV-2 clinical isolates, there were on average 9.8 (range 0-196, median 9) nucleotide changes and 5.6 (range 0-117, median 5) amino acid changes compared to the reference sequence (Supplementary Table 1). The average number of both nucleotide and amino acid changes increased steadily from December 2019 through September 2020 (Supplementary Table 1).

### 3.3 Genetic Changes in SARS-CoV-2 Replication Complex in Human Clinical Isolates

The number of amino acid substitutions within the RNA replication complex (nsp7, nsp8, nsp10, nsp12, nsp13, and nsp14) was calculated for the whole set of analyzed human clinical isolates. Low variation was observed across all genes encoding proteins of the RNA replication complex (see Supplementary Figures 1-6) with only 12 amino acid substitutions present in ≥0.5% of all clinical isolates (see Table 2). No amino acid substitutions with frequency ≥0.5% were observed in nsp8 or nsp10. The most prevalent substitution in RNA replication complex was nsp12 P323L, which was observed in 75,892/92,334 (82%) clinical isolates from 103 of 109 countries. Residue P323 of nsp12 is polymorphic across different coronaviruses, but conserved within each coronavirus (see Supplementary Figure 8 and Supplementary Table 3). Excluding P323L, no other substitutions were observed in 87% of SARS-CoV-2 clinical isolates in nsp12. No amino acid substitutions were observed in 97%, 97%, 99%, 88%, and 91% of clinical isolates in nsp7, nsp8, nsp14, nsp10, and nsp13, respectively.

**Table 2.** Amino Acid Sequence Variation observed in SARS-CoV-2 Replication Complex in Clinical Isolates.

| SARS-CoV-2 Gene | Isolates with No Amino Acid Substitutions Observed, % (N) | Average No of Amino Acid Substitutions Observed Per Isolate | Amino Acid Substitutions Observed in $\geq 0.5\%$ of Clinical Isolates | Frequency of Amino Acid Substitutions Observed in Clinical Isolates, % (N) |
| --- | --- | --- | --- | --- |
| Nsp12 | 15.1%<br>(13,978) | 0.96<br>(0-15) | P323L | 82.2% (75,892) |
|  |  |  | A97V | 1.6% (1,484) |
|  |  |  | T141I | 1.0% (899) |
|  |  |  | A449V | 0.6% (525) |
| Nsp7 | 97.3%<br>(89,810) | 0.03<br>(0-2) | S25L | 1.7% (1549) |
| Nsp8 | 97.3%<br>(89,879) | 0.03<br>(0-6) | - | - |
| Nsp14 | 98.8%<br>(91259) | 0.10<br>(0-14) | A320V | 0.6% (589) |
|  |  |  | F233L | 0.5% (460) |
|  |  |  | M501I | 0.5% (460) |
| Nsp10 | 87.9%<br>(81,142) | 0.01<br>(0-3) | - | - |
| Nsp13 | 91.4%<br>(84,368) | 0.16<br>(0-11) | Y541C | 2.4% (2,172) |
|  |  |  | P504L | 2.3% (2,148) |
|  |  |  | A18V | 0.8% (733) |
|  |  |  | E244D | 0.8% (693) |

### 3.4 Genetic Changes of SARS-CoV-2 Replication Complex in Mink Isolates

The frequency of amino acid substitutions within the RNA replication complex was assessed in N=333 mink isolates. Low variation was observed across all proteins with only N=12 amino acid substitutions in ≥0.5% of mink isolates. No amino acid substitutions with frequency ≥0.5% were observed in nsp14 or nsp10. Similar to human clinical isolates, the most prevalent substitution was nsp12 P323L, which was observed in 89.5% of mink isolates. Nsp12 T739I was observed in 26.1%, Nsp13 I285V in 27.9%, and Nsp13 R392C in 10.5% of mink isolates. Each of these substitutions except for Nsp12 P323L was observed in <0.5% of human clinical isolates (Table 3).

**Table 3.** SARS-CoV-2 Amino Acid Substitutions Observed in ≥0.5% Infected Mink Isolates.

| SARS-CoV-2 Gene | Minks with No Amino Acid Substitutions Observed, % (N) | Average No of Amino Acid Substitutions Observed Per Minks | Amino Acid Substitutions Observed in $\geq 0.5\%$ of Mink Isolates | Frequency Observed in Mink Isolates, % (N) | Frequency Observed in Clinical Isolates, % (N) |
| --- | --- | --- | --- | --- | --- |
| nsp12 | 8.1% (27) | 1.22 (0-3) | P323L | 89.5% (298) | 82.2% (75,892) |
|  |  |  | T739I | 26.1% (87) | <0.1% (28) |
|  |  |  | T803I | 1.8% (6) | 0.1% (88) |
|  |  |  | M196I | 1.2% (4) | <0.1% (17) |
|  |  |  | S913A | 0.9% (3) | <0.1% (2) |
|  |  |  | T293I | 0.6% (2) | <0.1% (25) |
|  |  |  | T85I | 0.6% (2) | <0.1% (58) |
| nsp7 | 98.8% (329) | 0.01 (0-1) | S25A | 0.6% (2) | <0.1% (2) |
| nsp8 | 99.4% (331) | <0.01 (0-1) | E23G | 0.6% (2) | <0.1% (1) |
| nsp14 | 98.8% (329) | 0.01 (0-1) | - | - | - |
| nsp10 | 100% (333) | 0 | - | - | - |
| nsp13 | 60.6% (202) | 0.41 (0-3) | I258V | 27.9% (93) | <0.1% (8) |
|  |  |  | R392C | 10.5% (35) | 0.3% (252) |
|  |  |  | A446D | 1.2% (4) | 0 |

### 3.5 Variant Frequencies Over Time in the SARS-CoV-2 RNA Replication Complex in Human Clinical Isolates

In the next step, we focused our analysis on temporal changes in genetic diversity of SARS-CoV-2 replication complex genes. In order to obtain reliable and interpretable data, we included only months with ≥1000 clinical isolate sequences available and focused on amino acid substitutions with total frequency ≥0.5% In the genes of the RNA replication complex, only the Nsp12 P323L substitution consistently increased in frequency in human clinical isolates over time. Nsp13 substitutions P504L and Y541C decreased in frequency over time. (Supplementary table 2). Available sequence data for mink isolates was limited and therefore excluded from this analysis.

### 3.6 Genetic Changes in SARS-CoV-2 Replication Complex in B.1.1.7 and B.1.351 Clinical Isolates

The nature and frequency of amino acid substitutions within the RNA replication complex was analyzed for human clinical isolates of the recently emerged B.1.1.7 and B.1.351 SARS-CoV-2 variants. Low variation was observed across all genes encoding proteins of the RNA replication complex in N=1987 B.1.1.7 and N=311 B.1.351 isolates, with only 5 and 10 amino acid substitutions present in ≥0.5% (see Tables 4 and 5), respectfully. The most prevalent substitution in RNA replication complex was nsp12 P323L, which was observed in 100% of the B.1.1.7 isolates and the B.1.351 isolates analyzed. Both SARS-Cov-2 variants also each contained one prevalent substitution in Nsp13; K460R was observed in 51% of B.1.1.7 isolates and T588I was observed in 14% of B.1.351 isolates. RDV has been shown to have similar antiviral activity against early-lineage SARS-CoV-2 and the B.1.1.7 variant in both human gastrointestinal and lung epithelial cells [26]. None of the other substitutions observed are expected to reduce RDV susceptibility, but experimental testing will be needed to validate.

**Table 4.** SARS-CoV-2 Amino Acid Substitutions Observed in >0.5% of B.1.1.7 Isolates.

| SARS-CoV-2 Gene | B.1.1.7 Isolates with No Amino Acid Substitutions Observed, % (N) | Average No of Amino Acid Substitutions Observed Per B.1.1.7 Isolate | Amino Acid Substitutions Observed in >0.5% B.1.1.7 Isolates | Frequency Observed in B.1.1.7 Isolates, % (N) | Frequency Observed in Expanded Clinical Isolate Set, % (N) |
| --- | --- | --- | --- | --- | --- |
| nsp12 | 0 | 1.06 (0-3) | P323L<br>P227L | 100% (1987)<br>2.7% (54) | 82.2% (75,892)<br><0.1% (91) |
| nsp7 | 99.1% (1969) | <0.01 (0-2) | - | - | - |
| nsp8 | 96.2% (1912) | 0.04 (0-2) | Q24R | 3.4% (67) | <0.1% (2) |
| nsp14 | 99.7% (1982) | <0.01 (0-1) | - | - | - |
| nsp10 | 99.2% (1972) | <0.01 (0-1) | - | - | - |
| nsp13 | 46.8% (930) | 0.54 (0-2) | K460R<br>P78S | 51.5% (1024)<br>0.7% (14) | <0.1% (37)<br><0.1% (24) |

**Table 5.** SARS-CoV-2 Amino Acid Substitutions Observed in >0.5% of B.1.351 Isolates.

| SARS-CoV-2 Gene | B.1.1.7 Isolates with No Amino Acid Substitutions Observed, % (N) | Average No of Amino Acid Substitutions Observed Per B.1.1.7 Isolate | Amino Acid Substitutions Observed in >0.5% B.1.1.7 Isolates | Frequency Observed in B.1.1.7 Isolates, % (N) | Frequency Observed in Expanded Clinical Isolate Set, % (N) |
| --- | --- | --- | --- | --- | --- |
| nsp12 | 0 | 1.05<br>(0-3) | P323L | 100% (311) | 82.2% (75,892) |
|  |  |  | D135Y | 1.0% (3) | <0.1% (2) |
|  |  |  | T85I | 0.6% (2) | <0.1% (58) |
|  |  |  | M124I | 0.6% (2) | <0.1% (33) |
| nsp7 | 99.3%<br>(309) | <0.01<br>(0-1) | - | - | - |
| nsp8 | 99.7%<br>(310) | <0.01<br>(0-1) | - | - | - |
| nsp14 | 99.7%<br>(1982) | <0.01<br>(0-1) | - | - | - |
| nsp10 | 98.1%<br>(305) | 0.02<br>(0-1) | S33N | 1.3% (4) | <0.1% (1) |
| nsp13 | 80.3%<br>(250) | 0.20<br>(0-2) | T588I | 13.2% (41) | <0.1% (58) |
|  |  |  | T351I | 1.3% (4) | <0.1% (26) |
|  |  |  | A598S | 1.0% (3) | <0.1% (73) |
|  |  |  | P47L | 0.6% (2) | 0.2% (229) |
|  |  |  | E261D | 0.6% (2) | 0.1% (101) |

### 3.7 Evaluation of Potential Pre-Existing Resistance to RDV in the SARS-CoV-2 Isolates

Amino acid sequence alignments demonstrate that nsp12 of SARS-CoV-2 exhibits 96% sequence identity with SARS-CoV, 71% with MERS-CoV, and 66% with MHV. However, the residues that define the nsp12 polymerase active site show significantly higher conservation, approaching 100% [17]. Furthermore, the nsp12 residues F476 and V553 that changed in MHV as a result of RDV in vitro selection in prior studies [19], corresponding to F480 and V557 in SARS-CoV-2, are 100% conserved across all Alpha, Beta, and Gamma CoVs (see Supplementary Figure 7). Finally, the nsp12 S861 residue that is responsible for RDV’s delayed chain termination is 100% conserved across Alpha, Beta, and Delta CoV’s, but is an alanine in Gamma CoV’s.

Among the whole set of >90,000 analyzed SARS-CoV-2 human clinical isolates, nsp12 amino acid substitution F480L was detected in a single isolate sequence and V557A in a single independent isolate sequence. This observation indicates an extremely low frequency (0.002%) of potentially pre-existing reduced susceptibility to RDV among circulating clinical isolates of SARS-CoV-2. S861F was observed in another single independent isolate sequence. In the B.1.1.7 and B.1.351 SARS-CoV-2 variants, no changes were observed at F480, V557, or S861. Independent studies assessing the phenotypic impact of nsp12 substitutions at F480, V557, and S861 residues in the context of SARS-CoV-2 viral replication are in progress and will be reported independently.

### 3.8 Evaluation of RdRp Residues Predicted to Impact RDV Susceptibility in the SARS-CoV-2 Clinical Isolates

All observed amino acid substitutions in nsp12, nsp7, nsp8, and nsp13 with a frequency greater than 0.5% in clinical isolates were mapped to a cryo-EM structure of the replication complex (PDB 6XEZ [4], see Figure 1). A similar map was generated on a homology model of nsp14 [27] (see Supplementary Figure 9). With respect to nsp12, none of the observed substitutions were determined to have any direct interaction with pre-incorporated RDV-TP, the site of delayed chain termination near S861, or the site of template dependent inhibition near A558. P323L, A97V and T141I are all solvent exposed residues which are > 25 Å from the polymerase active site. A449V, with an occurrence of 0.57%, is in the fingers domain and could have an indirect impact on residues in the F-motif, including V557 and A558. A similar indirect effect on RDV susceptibility was recently identified for Ebola virus, where an F548S mutation, also in the fingers domain, was seen to confer low level resistance [28]. While prediction of the potential effect of substitutions on drug resistance can involve multiple factors beyond a simple direct interaction with the inhibitor molecule, our assessment is that the potential for significant impact of the identified pre-existing nsp12 substitutions on RDV susceptibility appears to be low.

**Figure 1.**
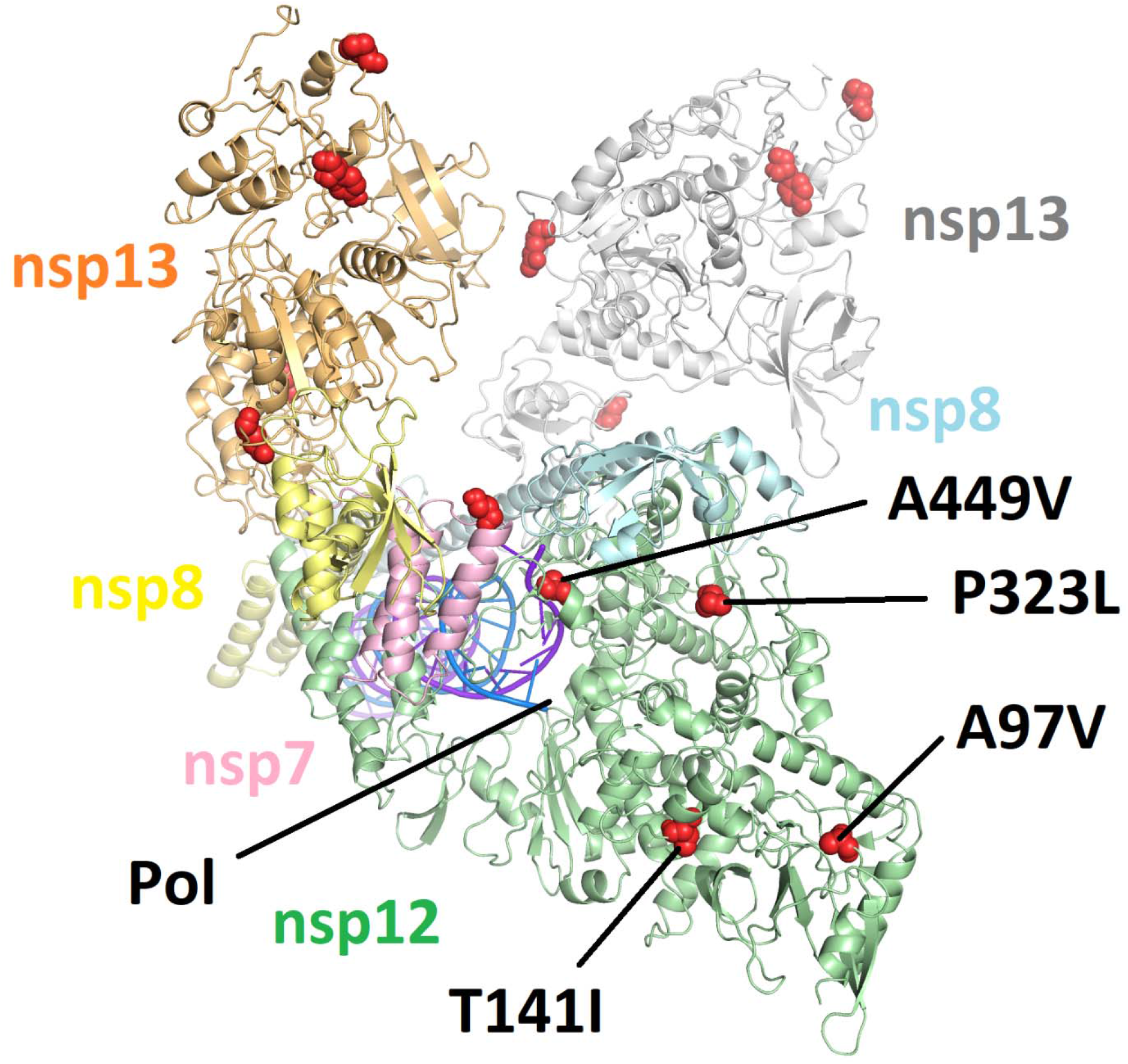
Amino acid substitutions (in red) with observed frequency > 0.5% as mapped to the most complete structure of the replication complex to date (PDB: 6XEZ) [4]. Nsp12 is in green, nsp7 is in pink, two subunits of nsp8 are in yellow and blue, and two subunits of nsp13 are in orange and white. Only amino acid substitutions in nsp12 are annotated. None of the mapped mutations are seen to interact directly with the nsp12 polymerase active site (Pol) or locations associated with either delayed chain termination or template dependent inhibition.

## 4. Discussion

Studies conducted early in the COVID-19 pandemic described minimal genetic variation between different SARS-CoV-2 clinical isolates [20]. Monitoring viral diversification in circulating SARS-CoV-2 isolates is important for identifying genetic changes that could potentially impact susceptibility to various COVID-19 therapeutics including the approved antiviral RDV. In the current study, the low genetic diversity in the RNA replication complex (nsp7, nsp8, nsp10, nsp12, nsp13, and nsp14) of SARS-CoV-2 was confirmed in a large set of global human clinical isolates. Nsp12 P323L is the most frequent substitution observed, increasing in frequency over time. Eleven other amino acid substitutions within the RNA replication complex occur in ≥0.5% of all isolates evaluated, but none have been increasing in frequency over time. From cryo-EM structures, none of the observed substitutions, including nsp12 P323L, were determined to have any direct interaction with pre-incorporated RDV-TP, the site of delayed chain termination, or the site of template dependent inhibition. Experimental validation of these finding is critical, are currently in progress, and will be reported as soon as available.

Additional datasets related to the recent SARS-CoV-2 variants identified in the United Kingdom and South Africa, B.1.1.7 and B.1.351, were analyzed for genetic changes in the RNA replication complex. Low variation was observed. Nsp12 P323L was observed in 100% of isolates from each lineage. Each SARS-CoV-2 variant also contained one nsp13 substitution, K460R in B.1.1.7 and T588I in B.1.351, that are unlikely to affect RDV susceptibility. RDV has been shown to have similar antiviral activity against early-lineage SARS-CoV-2 and the B.1.1.7 variant in both human gastrointestinal and lung epithelial cells [26].

With the recent transmission of SARS-CoV-2 to minks, genetic differences were evaluated for potential impact to RDV. In the mink dataset, low variation was also observed in the RNA replication complex and nsp12 P323L was the most frequent amino acid substitution, consistent with the human data. The transmission of SARS-CoV-2 from humans to other species and back to humans, may introduce further genetic diversification over time and should continue to be monitored for emergence of changes that could impact efficacy of therapeutics and vaccines. From the mink dataset evaluated in this report, the genetic diversification impacting RDV efficacy seems to be minimal.

Previous characterization of the in vitro resistance profile of RDV using related CoVs indicated a high genetic barrier to RDV resistance, and identified substitutions at two highly conserved nsp12 amino acid residues corresponding to F480 and V557 in SARS-CoV-2 to be associated with low-level reduced susceptibility of coronaviruses to RDV in vitro [19]. Among >90,000 circulating human clinical isolates of SARS-CoV-2 analyzed in the present study, only one isolate was found to have an amino acid substitution at residue F480, and one other independent isolate was found to have a substitution at residue V557. Another residue, S861, which was previously implicated in the delayed chain termination of RDV [17], was found to be conserved in all but one single isolate. In the B.1.1.7 and B.1.351 SARS-CoV-2 variants, no changes were observed at F480, V557, or S861.

Given the acute nature of SARS-CoV-2 infection, high sequence conservation in the SARS-CoV-2 RNA replication complex, and a short treatment duration with RDV, the probability of resistance development in patients treated with RDV is believed to be low. However, further studies are needed to assess the potential for resistance development specifically in the context of RDV treatment of patients with active SARS-CoV-2 infection. For instance, based on a recent study of Ebola virus resistance to RDV [28], focus on residues in the fingers domain, such as A449V observed in 0.57% of clinical isolates, and their potential impact on template dependent inhibition may be warranted. Currently, in vitro resistance selections with RDV and SARS-CoV-2 are being conducted to compare with the findings from previous studies with related CoVs [19] as well as potentially identifying additional residues that might confer reduced susceptibility to RDV. To determine the potential emergence of RDV resistance and its implications for a clinical response to RDV treatment, viral sequencing of SARS-CoV-2 isolates before, during, and after RDV treatment is in progress as a part of the analysis of both the completed and ongoing clinical studies. In addition, the SARS-CoV-2 sequence analysis methods combined with drug target structural analysis described herein can be applied to monitor pre-existing or emerging resistance to any other direct-acting antiviral drugs including other nucleoside RdRp inhibitors, viral protease inhibitors or neutralizing antibodies.

In conclusion, results of this extensive sequence analysis across >90,000 global SARS-CoV-2 isolates, including the recently emerged variants, highlights the low diversity and high genetic stability of the RNA replication complex, and suggest a minimal global risk of pre-existing SARS-CoV-2 resistance to RDV.

## Acknowledgements

We would like to thank the GISAID data submitters and curators supporting progress in the understanding of the SARS-CoV-2 genetics.

## Glossary

RdRp: RNA-dependent RNA Polymerase
CoV: Coronaviruses
SARS-CoV: Severe Acute Respiratory Syndrome Coronavirus
MERS-CoV: Middle Eastern Respiratory Syndrome Corona Virus
COVID-19: Coronavirus Disease 2019
RDV: Remdesivir
RDV-TP: Remdesivir-Triphosphate
MHV: Murine Hepatitis Virus

## Notes

Conflict of interest statement: Ross Martin, Jason Perry, Tomas Cihlar, Hongmei Mo, Danielle Porter, and Evguenia Svarovskaia are employees and stock holders of Gilead Sciences.

### Competing Interest Statement

All authors are employees and stock holders of Gilead Sciences.

### Summary of Updates

Updated to include analysis of UK and SA variant strains (section 3.6). Included analysis of observed variants in SARS-CoV-2 in %0.5% isolates in other coronaviruses and isolates from human coronavirus infections (supplementary). Added reference for experimental findings of RDV in original and B.1.1.7 strain.

