## supplemental for "Genetic Conservation of SARS-CoV-2 RNA Replication Complex in Globally Circulating Isolates and Recently Emerged Variants from Humans and Minks Suggests Minimal Pre-Existing Resistance to Remdesivir"

**Supplementary Table 1. Number of Changes in the SARS-CoV-2 Clinical Isolate Sequences**

|  | Number of SARS‑CoV-2 clinical isolate sequences analyzed | Average number of nucleotide changes per full genome | Percent of clinical isolates with no nucleotide changes observed | Average number of amino acid changes across all open reading frames | Percent of clinical isolates with no amino acid changes observed |
| --- | --- | --- | --- | --- | --- |
| December 2019 | 18 | 2.9 | 33.3% | 1.4 | 55.5% |
| January 2020 | 452 | 3.3 | 12.4% | 1.9 | 19.4% |
| February 2020 | 1,134 | 4.8 | 2.6% | 3.0 | 5.6% |
| March 2020 | 29,650 | 7.5 | 0.3% | 4.4 | 0.8% |
| April 2020 | 27,118 | 8.9 | 0 | 5.2 | 0.1% |
| May 2020 | 11,834 | 10.7 | 0 | 6.2 | 0.1% |
| June 2020 | 7,928 | 12.2 | 0 | 6.8 | 0 |
| July 2020 | 7,932 | 14.2 | 0 | 7.4 | 0 |
| August 2020 | 5,026 | 15.8 | 0 | 8.8 | 0 |
| September 2020 | 301 | 16.2 | 0 | 8.8 | 0 |
| Total^a^ | 92,344 | 9.8 | 0.2% | 5.6 | 0.5% |

^a^ n=941 clinical isolates without month of collection provided

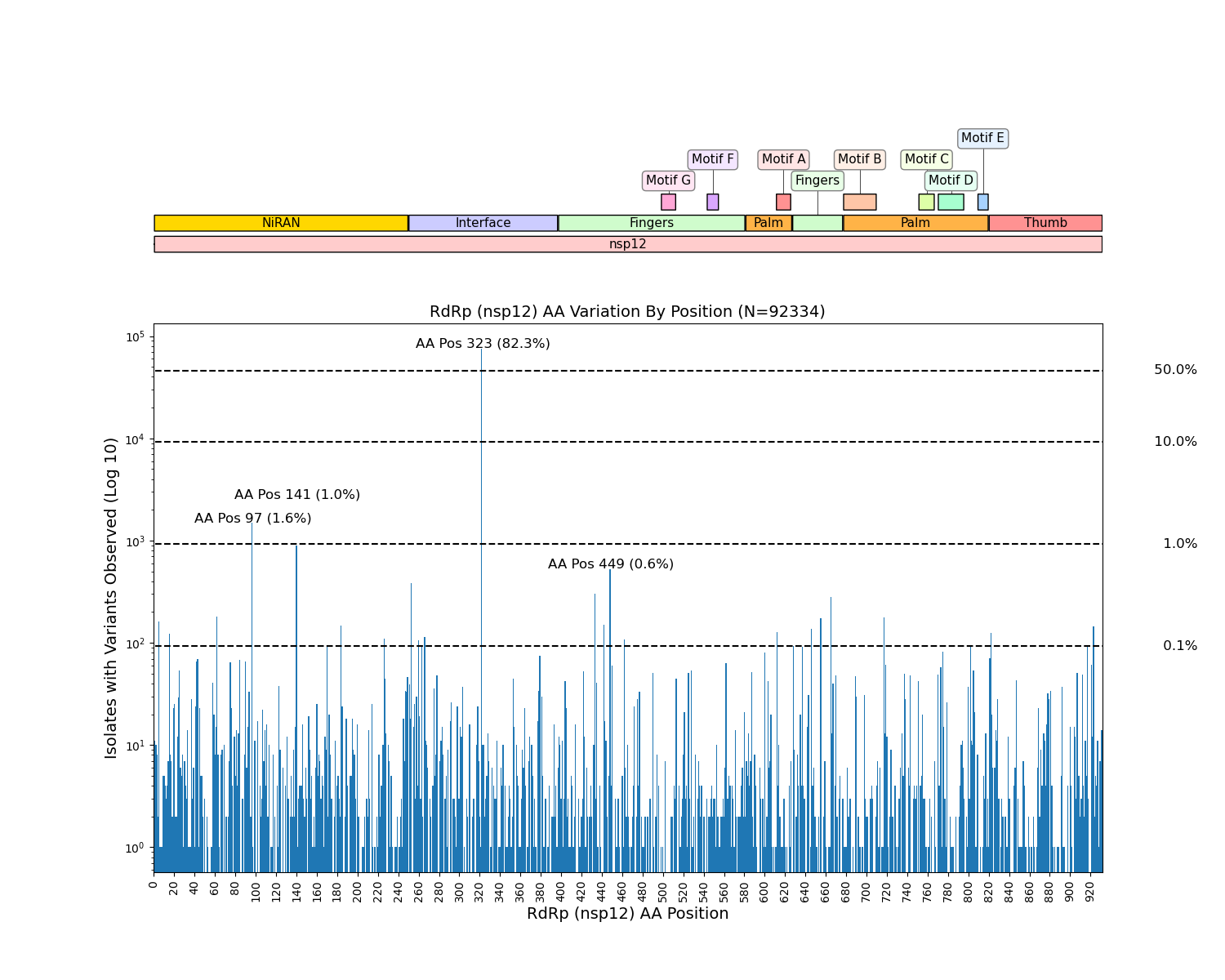

**Supplementary Figure 1. Amino Acid Variation By Position in SARS-CoV-2 RdRp (nsp12)**

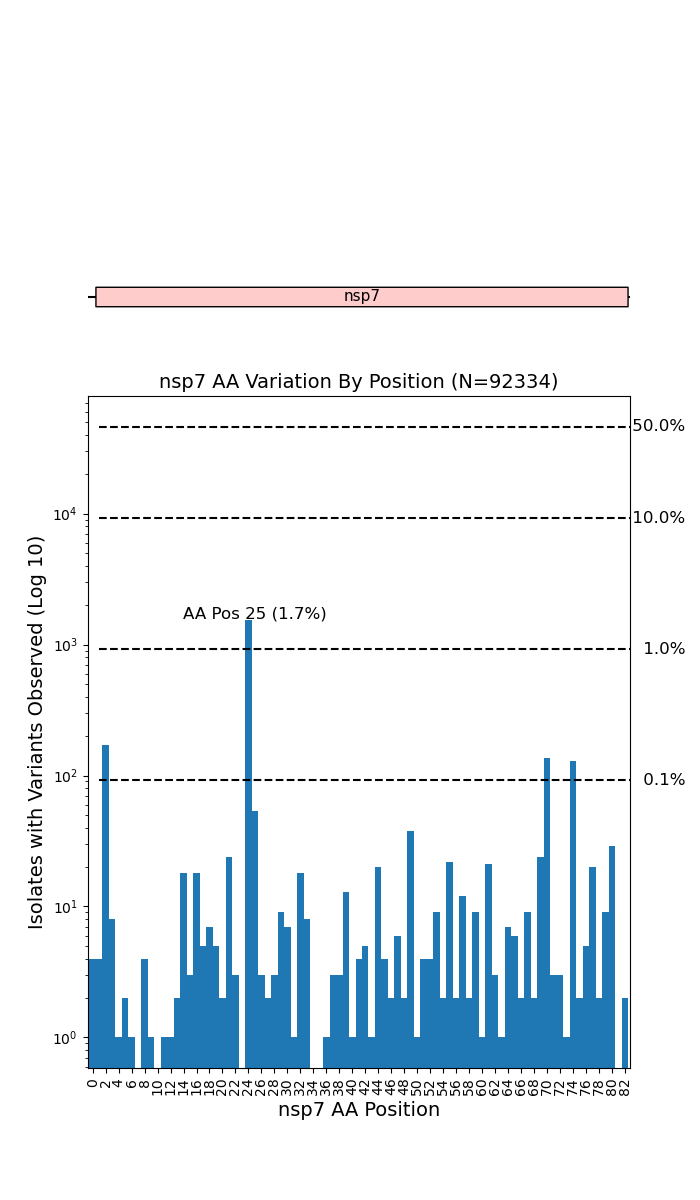

**Supplementary Figure 2: Amino Acid Variation By Position in Nsp7**

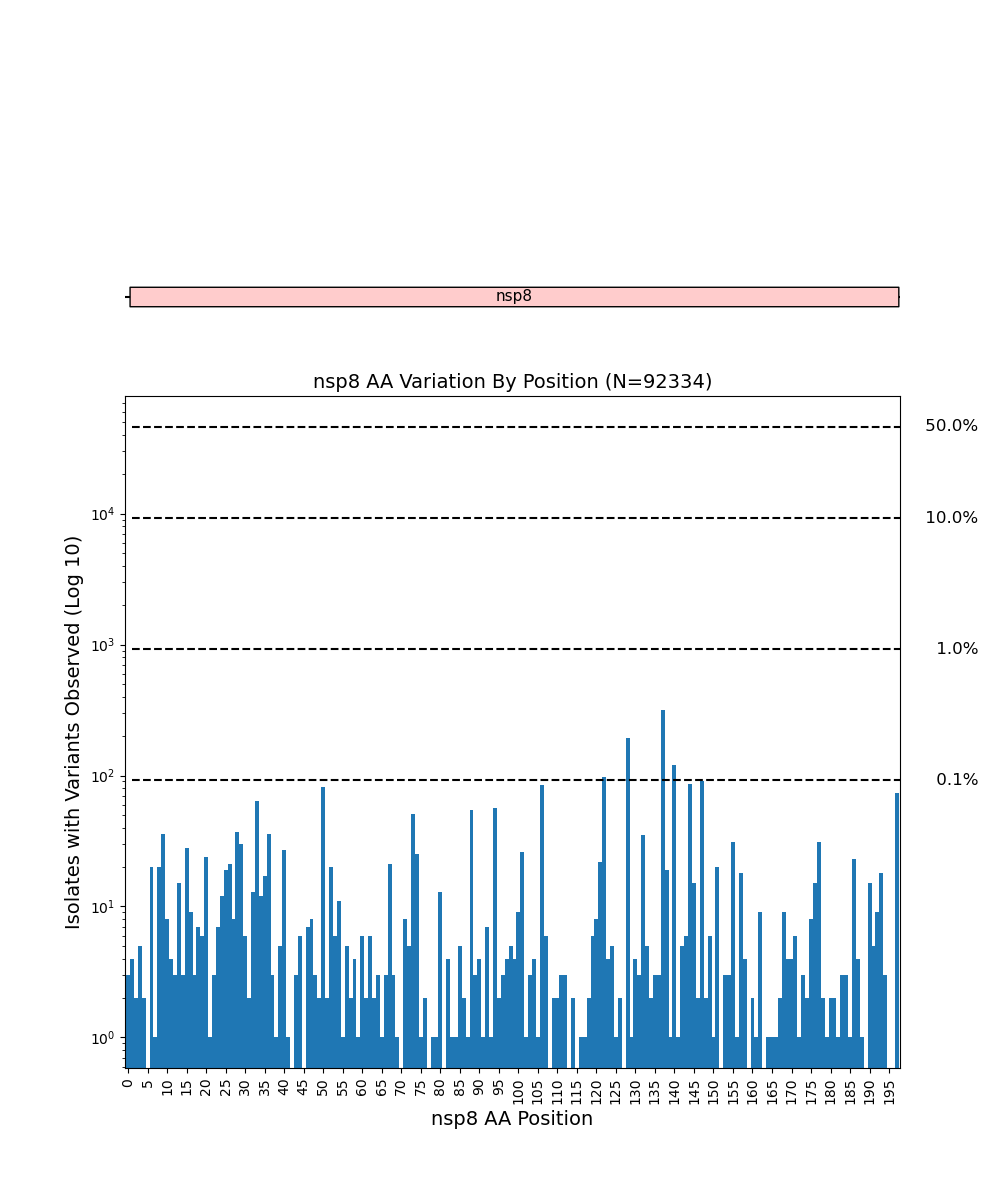

**Supplementary Figure 3: Amino Acid Variation By Position in Nsp8**

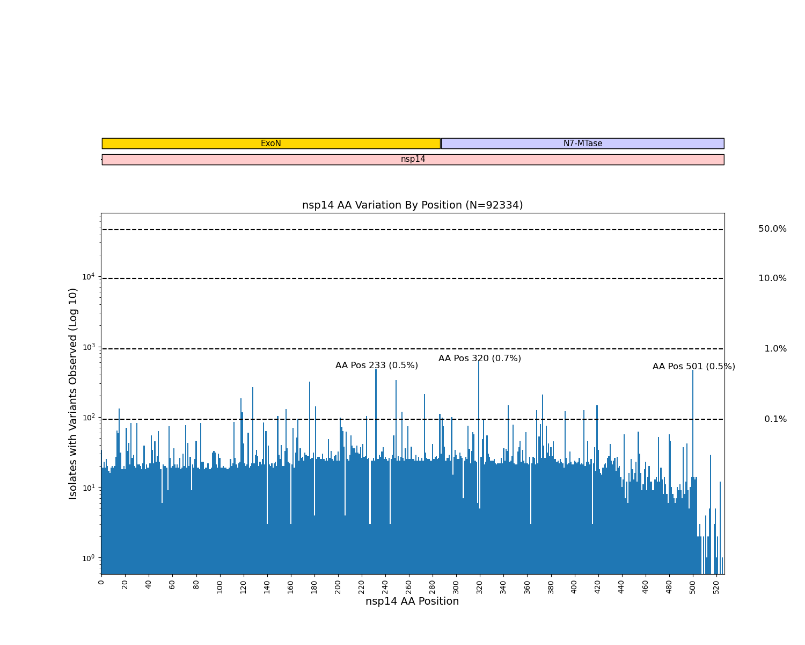

**Supplementary Figure 4: Amino Acid Variation By Position in Nsp14 Exonuclease**

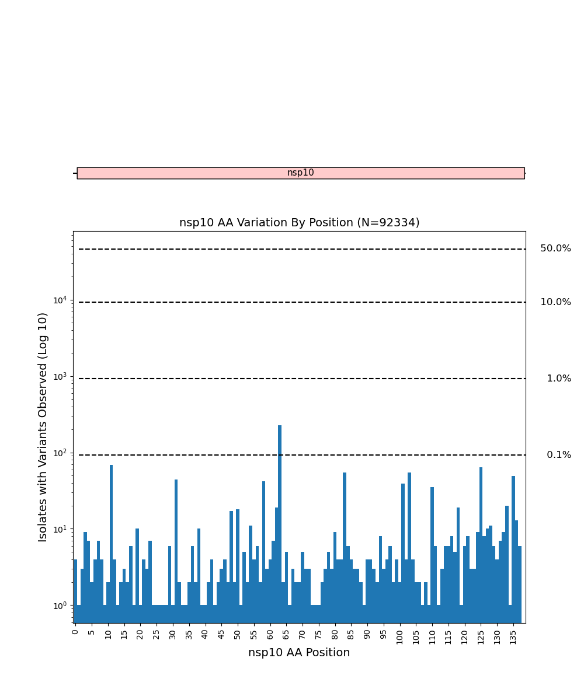

**Supplementary Figure 5: Amino Acid Variation By Position in Nsp10**

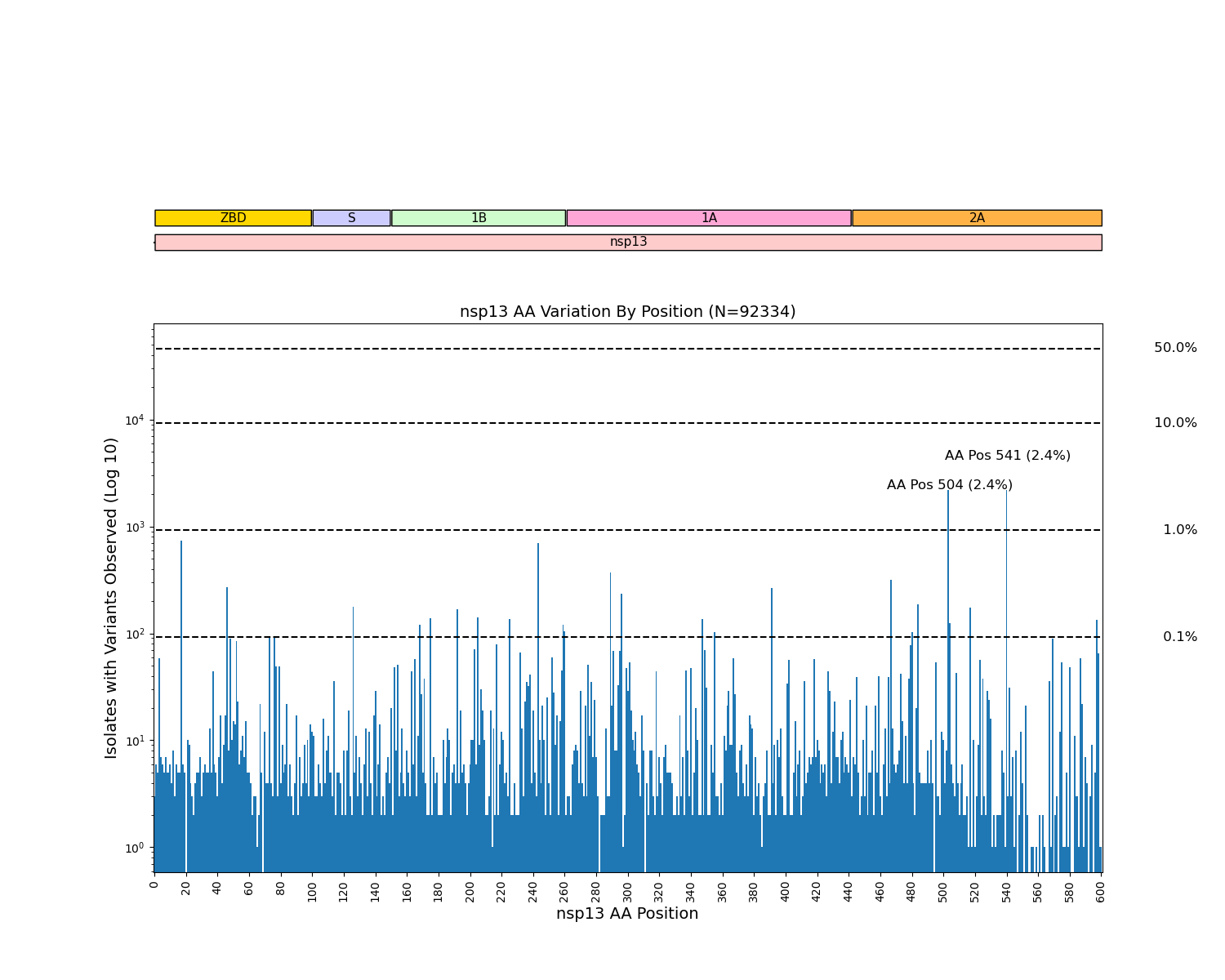

**Supplementary Figure 6: Amino Acid Variation By Position in Nsp13 Helicase**

**Supplementary Table 2 Change in Frequency Over Time of Amino Acid Substitutions in SARS-CoV-2 RNA Replication Machinery Observed in ≥0.5% of Clinical Isolates**

|  | Clinical Isolate Count (N) | 92,344 | 1,134 | 29,650 | 27,118 | 11,834 | 7,927 | 7,932 | 5,026 |  |  |
| --- | --- | --- | --- | --- | --- | --- | --- | --- | --- | --- | --- |
| SARS-COV-2 Gene | Amino Acid Substitution | Total Frequency (%) | February Frequency (%) | March Frequency (%) | April Frequency (%) | May Frequency (%) | June Frequency (%) | July  Frequency (%) | August Frequency (%) | Slope of Regression Line | p-value |
| nsp12 | P323L | 82.18 | 19.49 | 67.95 | 84.54 | 93.67 | 96.22 | 98.5 | 98.57 | 11.07 | 0.02 |
| nsp13 | P504L | 2.33 | 4.59 | 4.85 | 1.71 | 0.9 | 1.01 | 0.03 | 0.02 | -0.86 | <0.01 |
| nsp13 | Y541C | 2.35 | 4.5 | 4.97 | 1.74 | 0.87 | 0.83 | 0.03 | 0.02 | -0.87 | <0.01 |

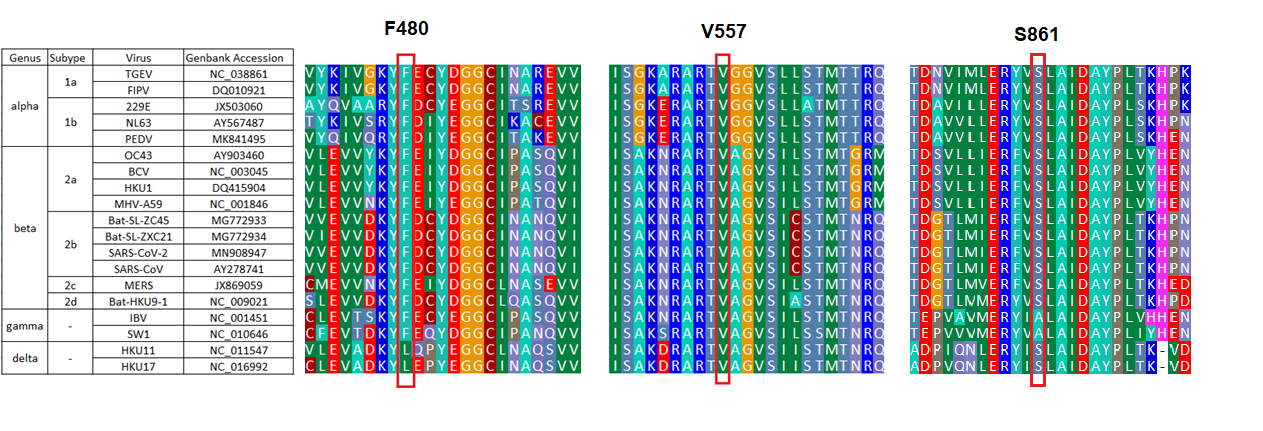

**Supplementary Figure 7**. Conservation of nsp12 amino acid residues in predicted resistance residues to RDV (F480, V557) and the residue responsible for RDV’s delayed chain termination (S861) across coronaviruses. Amino acid alignment of nsp12 includes Alpha coronaviruses TGEV, FIPV, 229E, NL63, PEDV; Beta coronaviruses OC43, BCV, HKU1, MHV-A59, Bat-SL-ZC45, Bat-SL-ZXC21, SARS-CoV-2, SARS-CoV, MERS, Bat-HKU9-1; Gamma coronaviruses IBV, SW1; Delta coronaviruses HKU11, HKU17.

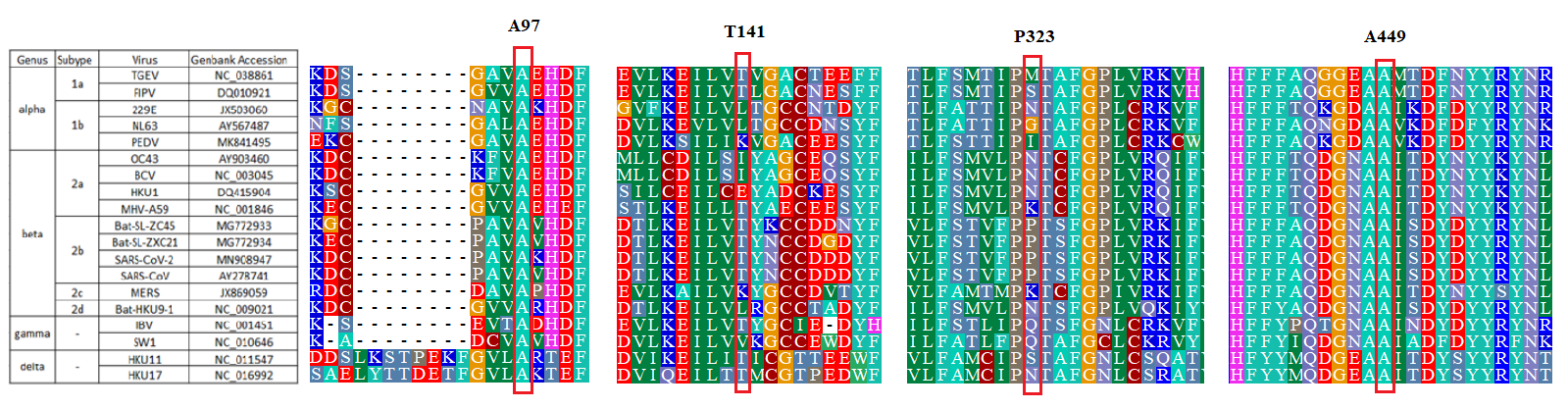

**Supplementary Figure 8**. Conservation across coronaviruses of nsp12 amino acid residues with substitutions observed in >0.5% of SARS-CoV-2 human clinical isolates. Amino acid alignment of nsp12 includes Alpha coronaviruses TGEV, FIPV, 229E, NL63, PEDV; Beta coronaviruses OC43, BCV, HKU1, MHV-A59, Bat-SL-ZC45, Bat-SL-ZXC21, SARS-CoV-2, SARS-CoV, MERS, Bat-HKU9-1; Gamma coronaviruses IBV, SW1; Delta coronaviruses HKU11, HKU17.

**Supplementary Table 3**. Observed Amino Acids in Other Coronaviruses from Human Infections at nsp12 Positions with Substitutions Observed in >0.5% in SARS-CoV-2 Human Clinical and Mink Isolates

| Host of Observed  SARS-CoV-2 AA | SARS-CoV-2 nsp12 AA Position | SARS-CoV-2 Observed Amino Acids  (N=92344) | 229E Observed Amino Acids  (N = 32) | NL63 Observed Amino Acids  (N = 58) | HKU1 Observed Amino Acids  (N = 25) | OC43  Observed Amino Acids  (N = 137) | SARS1  Observed Amino Acids  (N = 59) | MERS Observed Amino Acids  (N = 336) |
| --- | --- | --- | --- | --- | --- | --- | --- | --- |
| human | 97 | A (98.4%)  V (1.6%) | A (100%) | A (100%) | A (100%) | A (100%) | A (100%) | A (100%) |
|  | 141 | T (99%)  I (1.0%) | V (100%) | V (100%) | I (84%)  V (16%) | L (100%) | T (100%) | V (100%) |
|  | 323 | L (82.2%)  P (17.8%) | N (100%) | G (100%) | N (100%) | N (100%) | P (100%) | K (100%) |
|  | 449 | A (99.4%)  V (0.6%) | A (100%) | A (100%) | A (100%) | A (100%) | A (100%) | A (100%) |
| mink | 85 | T (99.4%)  I (0.6%) | S (100%) | S (100%) | T (96%)  N (4%) | K (98.5%)  E (1.5%) | T (100%) | H (100%) |
|  | 196 | M (98.8%)  I (1.2%) | M (100%) | M (100%) | L (100%) | L (100%) | M (100%) | M (99.4%)  I (0.6%) |
|  | 293 | T (99.4%)  I (0.6%) | D (100%) | D (100%) | K (100%) | P (100%) | T (100%) | T (100%) |
|  | 739 | T (73.9%)  I (26.1%) | E (100%) | E (100%) | Y (100%) | S (100%) | H (100%) | P (100%) |
|  | 803 | T (99.4%)  I (0.6%) | E (100%) | E (100%) | N (100%) | H (97.8%)  N (1.4%)  Y (0.8%) | T (100%) | T (100%) |
|  | 913 | S (99.1%)  A (0.9%) | S (93.7%)  F (6.3%) | D (100%) | L (100%) | Q (100%) | S (100%) | A (100%) |

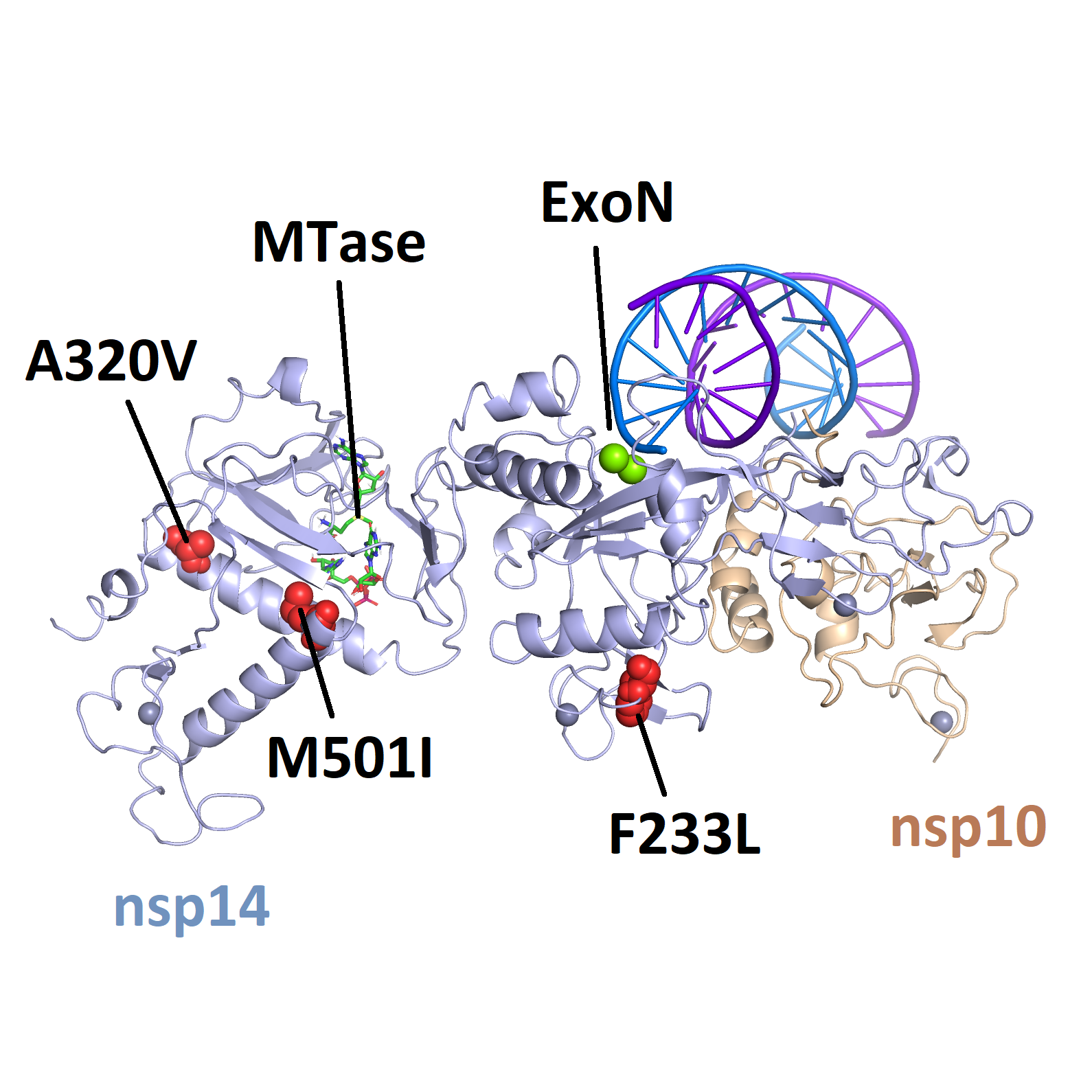

**Supplementary Figure 9.** Location of amino acid substitutions with >0.5% frequency as mapped onto a structure of nsp10/nsp14. The structure is a homology model based on an x-ray crystal structure of the SARS-CoV complex (PDB 5NFY) [23]. dsRNA has been modeled into the exonuclease (ExoN) active site for reference. The *N7*-methyltransferase (MTase) site is also indicated.
